# Mitag4taxa: Extracting SSU rRNA Illumina reads from metagenomes for taxonomic classification

**DOI:** 10.64898/2026.05.01.722230

**Authors:** Yinghui He, Yiling Du, Loi Nguyen, Yong Wang

## Abstract

The prevailing taxonomic profiling methods for an environmental sample rely heavily on PCR amplification of SSU ribosomal RNA (rRNA) genes and genome-based reference databases. Identification and extraction of Illumina metagenomics sequencing data are PCR independent but technically challenging in recognition of the SSU rRNA fragments. Here we present Mitag4taxa, a computational pipeline designed for taxonomic profiling of microbial communities from metagenomic Illumina sequencing reads containing rRNA tags (mitag). A Hidden Markov Model (HMM) of SSU rRNA genes and those for the V4 region of 16S rRNA and the V9 region of 18S rRNA genes were created, respectively, using the representative sequences of different families and corresponding hypervariable regions in the SILVA database. The pipeline identifies and extracts 16S and 18S rRNA gene fragments along with the quality score from metagenomic or metatranscriptomic datasets using HMM search integrated with the models. The hypervariable regions, including the V4 region of 16S rRNA and the V9 region of 18S rRNA genes, can be further scanned and recruited for taxonomic classification and biodiversity estimate. To demonstrate its high reliability, the performance of Mitag4taxa was evaluated using both real and simulated datasets. In human gut metagenomic assessments, taxonomic profiles derived from Mitag4taxa showed high consistency with those based on conventional 16S rRNA gene amplicons, identifying dominant families such as *Bacteroidaceae* and *Prevotellaceae* with similar relative abundances. Statistical analyses confirmed highly significant positive correlations between Mitag4taxa and amplicon-based community structures. The 18S V9 module was further validated using shotgun metagenomic data from deep-sea sediment cores, successfully recovering key eukaryotic taxa such as *Collodaria* and *Leotiomycetes*. Furthermore, benchmarking against the RiboTagger software using CAMI marine simulated datasets revealed that Mitag4taxa achieved a higher average F1 score and lower error metrics. Overall, Mitag4taxa provides a complementary rRNA gene amplicon- and genome-independent strategy for microbial community profiling, enabling improved detection of both prokaryotic and eukaryotic taxa from metagenomic and metatranscriptomic sequencing data.

## Introduction

High-throughput sequencing technologies have greatly advanced the understanding of microbial diversity and function across various environments including gut, soil and marine ecosystems^1,2^. Copy numbers of biomarkers such as small subunit (SSU) ribosomal RNA (rRNA) genes and ribosomal protein-coding genes in the environmental DNA are counted for detection of microbial community structure of a sample. PCR amplification of SSU rRNA genes for sequencing and quantification of the gene copies remains the most widely used method for profiling microbial taxonomic composition. However, it provides trivial information about the functional capabilities of microorganisms regardless the amplification biases due to primer selection^3^. With more rare microbial phyla from extreme ecological niches being discovered, designation and selection of PCR primers that optimally cover all the taxa in an environmental sample become highly challenging. In contrast, PCR-independent metagenomic sequencing provides an alternative approach to microbial community structure examination, in addition to exploration of microbial metabolic functions and activity dynamics. Identification and recruitment of SSU rRNA genes and ribosomal protein-coding genes from the raw Illumina sequencing data are more straightforward to disclose the community structure. With a high sequencing depth, thousands of >100 bp reads containing the biomarker information are sufficient to accurate taxonomic assignment of the reads^4^. However, the short reads cannot contain the full length of the SSU rRNA genes. As a result, the reads from different hypervariable regions cannot be aligned well for subsequent biodiversity estimate and similarity analysis of the community structures. To achieve better performance of the biomarker reads, we need to specify the hypervariable region to V4 of 16S rRNA genes^5^ and V9 of 18S rRNA^6^ genes for analysis of biodiversity and community structural similarity.

At present, the metagenomes have been applied for indirect prediction of the taxonomic composition through similarity search of the reads in the genome databases that are associated with taxonomic information of the genomic sequences^7^. The metagenome assembled genomes (MAGs) also harbor biomarkers for taxonomic assignment, although regarded as draft genomes with various completeness levels. Using coverage level of the MAGs by the reads, the relative abundance of the taxa represented by the MAGs can be estimated. The read recruitment percentages by all the MAGs are indicators of the community structures of a sample. However, the low-abundance rare species is always not deeply sequenced in a metagenome, which results in lack of their MAGs for taxonomic profiling. Nearly half of the sequencing reads cannot be assigned to the draft genomes, indicating a low efficiency of this method for reconstruction of the microbial community structure^8^.

Several computational approaches have been developed to identify the taxonomic composition and relative abundance of microorganisms directly from metagenomic reads. Kraken2 classifies sequencing reads by breaking them into smaller fragments known as k-mers, which are then mapped to the lowest common ancestor in a pre-constructed reference database through exact alignment^9^. When used in combination with Bracken^10^, Kraken2 further estimates the relative abundance of taxa. However, the classification accuracy of Kraken2 strongly depends on the completeness and quality of its reference database. The standard Kraken2 database, which includes bacterial, archaeal, and viral genomes, exceeds 100 Gbp, requiring substantial computational and storage resources. MetaPhlAn adopts a marker gene–based strategy^11^. It identifies taxonomic composition by aligning metagenomic or metatranscriptomic reads against a database of clade-specific marker genes, typically single-copy genes unique to each taxon^11^. Because it only targets a subset of representative marker genes rather than entire genomes, MetaPhlAn is both computationally efficient and storage-friendly compared to Kraken2. However, the completeness and accuracy of MetaPhlAn’s database depend heavily on the availability of fully sequenced genomes and high-quality MAGs. Although the number of reference genomes and MAGs has rapidly increased in recent years, genomic information for microorganisms inhabiting specialized or extreme environments, such as marine or polar ecosystems, remains relatively limited^12^. This scarcity of reference sequences constrains the classification accuracy of MetaPhlAn and similar methods when applied to such environments.

In this study, an efficient and lightweight solution for identifying microbial community composition and diversity from metagenomic datasets is developed, particularly for users with limited computational resources. This method identifies and utilizes SSU rRNA fragments, the V4-region reads of the 16S rRNA genes and the V9-region reads of the 18S rRNA gene to infer taxonomic composition directly from metagenomic sequences^13,14^. Therefore, we name it as Mitag4taxa. The pipeline first identifies these SSU rRNA sequences through hidden Markov model (HMM) ^15^ and then extract them for further scanning of the V4 and V9 hypervariable regions, followed by taxonomic assignment and biodiversity estimate. This strategy skips bias-introducing PCR amplifications and effectively reduces the need for large reference databases during the miTag identification. By leveraging the universally variable nature of the SSU rRNA genes, Mitag4taxa enables comprehensive and comparable microbial profiling across diverse environments, including those where reference genomes remain scarce. Furthermore, Mitag4taxa provides a practical framework for microbial community characterization, supporting ecological and functional studies even in resource-constrained research settings.

## Methods

### Construction of Region-Specific HMMs

Taxonomic reference sequences for the large subunit (LSU) and SSU rRNA sequences were retrieved from the SILVA database (release 138.1)^16^, utilizing the 99% non-redundant (NR99) datasets in the aligned format^16^. These reference sequences were further subdivided into three domains (Archaea, Bacteria, and Eukaryota) based on the taxonomic annotation of each sequence by the SILVA database. The aligned sequences were then individually converted to Stockholm format and used to construct hidden Markov models (HMMs) with the hmmbuild program from the HMMER package^15^.

To create the HMM models for identification the specific SSU variable regions from the Illumina reads, we curated the sub-regions of the representative SSU sequences in alignment format (Figure 1A). Firstly, the 16S rRNA genes were retrieved from the pre-aligned SSU sequences of the SILVA 138 database. To maintain phylogenetic diversity, a representative sequence was randomly sampled from each of ten randomly-selected, distinct families within each order. If there are less than 10 families in an order, each family has a representative sequence in the curated dataset. In case that a phylum does not have sequences at order level, ten sequences were randomly recruited from the classes into the dataset. The V4 region was extracted by an in-house python program using the locations determined by mapping the 515F/806R primer set (515F: GTGCCAGCMGCCGCGGTAA; 806R: GGACTACVSGGGTATCTAAT)^3^ onto the corresponding alignment. Secondly, eukaryotic 18S rRNA genes were downloaded from the PR2 database^17^, which collects more sequences that SILVA. A full-length 18S rRNA sequence of *Guillardia theta*, retrieved from NCBI, was inserted as a positional anchor and aligned together with these reference PR2 sequences using MAFFT^18^. Universal V9 primers (1380F: CCGGTGAATATTGVGYGCAA; 1510R: GGTTACCTTGTTACGACTT)^19^ were mapped onto the aligned *Guillardia theta* sequence to determine the corresponding alignment coordinates of the V9 region. The homologous V9 segments from all the aligned sequences were extracted by the same in-house program. For the 18S V9 regions, representative sequences were randomly selected to ensure phylogenetic diversity as aforementioned. The 16S V4-specific and 18S V9-specific HMMs were then created with the hmmbuild program from the HMMER package^15^.

**Figure 1.**
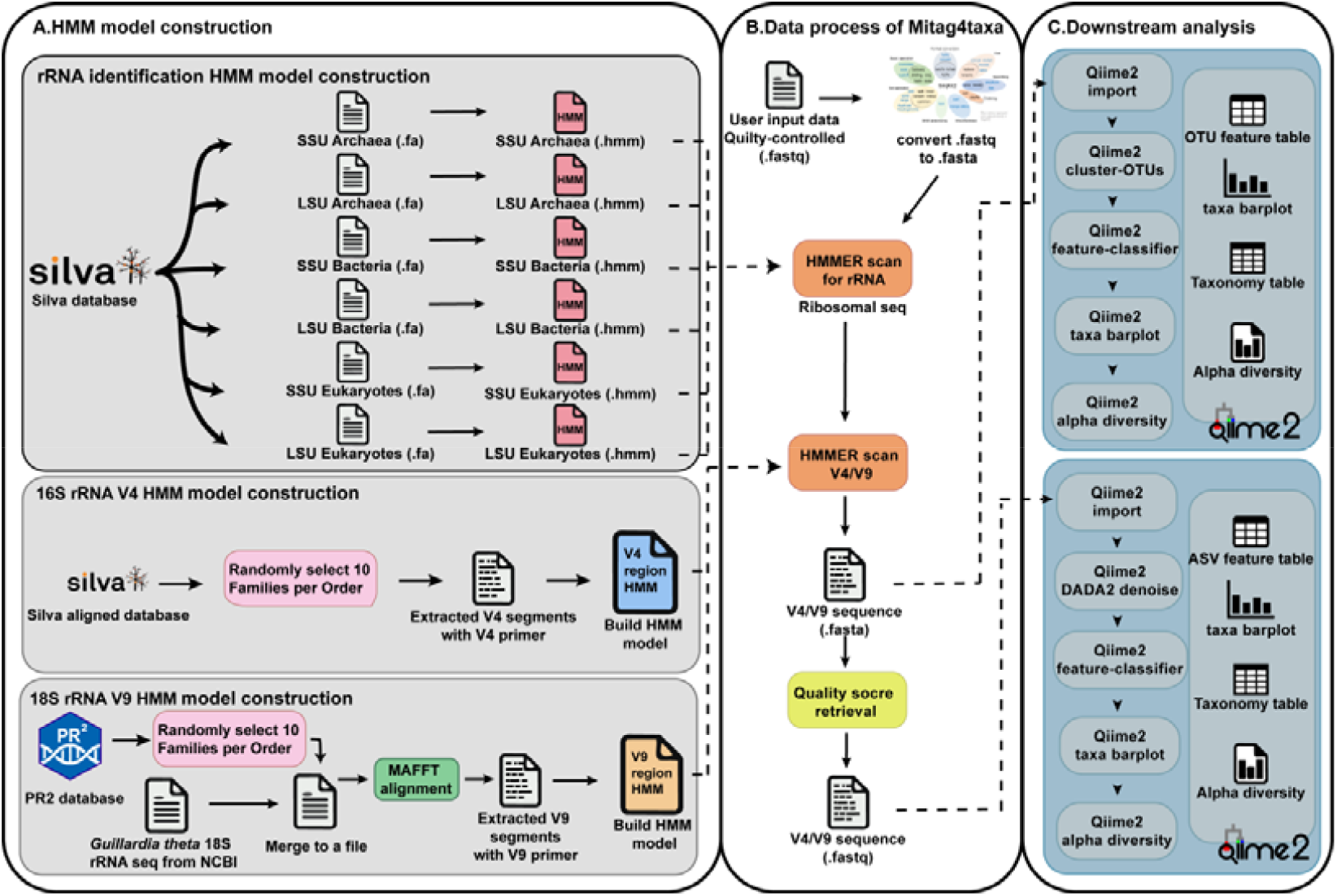
Schematic illustration of the Mitag4taxa software workflow. A. Hidden Markov Model (HMM) model construction module illustrates the steps for construction of HMM models. B. The core data processing of Mitag4taxa. Quality-controlled FASTQ files are converted to FASTA format for HMMER-based rRNA scanning. C. Downstream analysis is integration with QIIME2. The extracted sequences are processed through two parallel paths: OTU clustering or ASV-based denoising (DADA2).

### Mitag extraction and taxonomic classification

After the quality control with fastq^20^, the Illumina sequencing reads in FASTQ format are transferred to files without quality value (in FASTA format) with seqkit (Figure 1B)^21^. Subsequently, the previously constructed SSU and LSU HMM models for domains were employed to identify ribosomal RNA gene sequences from the query data (Figure 1B). The 16S and 18S rRNA reads in the SSU dataset were scanned against the corresponding region-specific HMM profiles using HMMER, and the aligned segments matching the V4 or V9 models were delineated based on HMM alignment coordinates, thereby yielding the final V4 and V9 fragments for downstream taxonomic analysis. The extracted V4 and V9 sequences are stored in a file in FASTA format, and the corresponding quality scores of these sequences are retrieved from the original Illumina sequencing data in FASTQ format to construct a new file. Therefore, two types of 16S V4 and 18S V9 sequences are obtained: one in FASTA format without quality scores, and the other in FASTQ format containing both sequences and corresponding quality scores.

For the FASTA sequences, the files are imported into the Qiime2^22^ to cluster the reads into operational taxonomic units (OTUs) at a 99% similarity threshold with cluster-OTUs module. For the sequences in FASTQ format, Illumina sequencing errors were corrected using DADA2^23^. Subsequent taxonomic assignment for both OTUs and ASVs was executed using the classify-sklearn plugin against the SILVA 138 (for prokaryotes) or PR2 (for eukaryotes) reference databases^17^. Alpha diversity indices, including Chao1, Shannon, and Simpson, were calculated using the alpha-rarefaction function in QIIME2.

### Software evaluation

To evaluate the performance of Mitag4taxa in identifying microbial community composition, we used both real and simulated datasets. For the real-data assessment, we analyzed a publicly available human gut metagenomic dataset (BioProject accession number: PRJNA725613)^24^. This dataset for 16 fecal samples collected from hospitalized patients were generated by a hospital in central China. The patients were classified into three groups: Group A (adenoma), Group C (colorectal cancer), and Group H (healthy controls). Each sample was sequenced using two methods: amplicon sequencing and whole metagenome sequencing. Details of DNA extraction, amplicon library preparation, and metagenomic sequencing procedures are provided in the original publication^24^. For the amplicon sequencing data, we obtained the V4 region sequences of the 16S rRNA gene, provided in both quality-preserved (.fastq) and quality-filtered (.fasta) formats. The prokaryotic community structures were reconstructed using two standard approaches: DADA2 (denoising-based amplicon variant inference) and OTU clustering. The community composition and microbial diversity inferred from these amplicon-based analyses were then compared to that obtained from the metagenomic data processed by Mitag4taxa, allowing us to assess the consistency and accuracy of taxonomic identification between methods.

To rigorously evaluate the methodological performance of Mitag4taxa for 18S-based taxonomic profiling, we analyzed a publicly available shotgun metagenomic dataset deposited under BioProject accession number PRJNA766251^25^. This dataset comprises Illumina whole-metagenome sequencing reads derived from Santa Barbara Basin deep-sea sediment cores, thereby providing an appropriate benchmark for assessing marker-gene recovery directly from metagenomic data. Raw paired-end FASTQ files were processed using Mitag4taxa with the 18S rRNA gene specified as the target marker. The pipeline first performed rRNA gene prediction to identify candidate SSU rRNA fragments from the total metagenomic reads. Subsequently, predicted 18S sequences were extracted, and the V9 hypervariable region was retrieved for downstream taxonomic classification.

To further assess the taxonomic classification accuracy of Mitag4taxa, we compared its performance with that of RiboTagger^26^ using nine simulated marine datasets from the second round of the CAMI (Critical Assessment of Metagenome Interpretation) challenge^27^. RiboTagger was selected for comparison due to its use of a similar approach, which involves extracting the V4 region to estimate relative abundances in prokaryotic metagenomic and metatranscriptomic datasets. We calculated the relative abundance in the datasets using the Mitag4taxa and RiboTagger at genus level, and transfer the output file into the CAMI format file. These CAMI-format profiles were then submitted to the official CAMI evaluation platform (https://www.cami-challenge.org/submit/), where software performance was quantified by calculating the F1 score, weighted UniFrac error, Bray–Curtis distance, and L1 norm error against the gold standard profiles.

To evaluate the computational efficiency of Mitag4taxa, a runtime benchmarking analysis comparing it against three widely used taxonomic profiling tools: Kraken2, MetaPhlAn, and RiboTagger was conducted. The benchmarking was performed using six human gut metagenomic samples (A1–A6, derived from BioProject PRJNA725613) to evaluate performance on high-complexity biological datasets. To ensure experimental rigor and comparability, all tools were executed on an identical hardware configuration with a fixed resource allocation of 2 CPU cores.

## Results and discussion

### Evaluation of Mitag4taxa using real testing data

We validated the performance of Mitag4taxa using real public metagenomic data and compared the taxonomic profiling results with those derived from conventional 16S rRNA amplicon sequencing of the same samples. This analysis was designated to evaluate the consistency and reliability of Mitag4taxa in detecting microbial community composition across different strategies. At the family level, the most abundant taxa identified from metagenomic data analyzed with Mitag4taxa using the OTU-based approach were *Bacteroidaceae, Prevotellaceae*, and *Lachnospiraceae*, with average relative abundances of 30.82 ± 6.00%, 20.64 ± 7.33%, and 11.83 ± 2.38% (mean ± sem), respectively (Figure 2a, Table S1). When applying the ASV-based method within Mitag4taxa to the same dataset, the top-ranked families remained *Bacteroidaceae, Prevotellaceae*, and *Lachnospiraceae*, showing relative abundance of 32.57 ± 6.93%, 25.43 ± 8.55%, and 10.12 ± 2.10%, respectively, highly consistent with those based on the OTU methods (Figure 2b, Table S2). A similar taxonomic pattern was observed in the 16S rRNA amplicon sequencing data, where *Bacteroidaceae, Prevotellaceae*, and *Lachnospiraceae* were again the dominant families, accounting for 28.28 ± 5.36%, 20.57 ± 6.38%, and 13.54 ± 2.54%, respectively (Figure 2c, Table S3). Mantel tests were employed to evaluate the congruence of community structures derived from amplicon, ASV-based Mitag4taxa, and OTU-based Mitag4taxa pipelines. All pairwise comparisons revealed highly significant positive correlations (*P* = 0.001, Table 1).

**Table 1.** Mantel test results comparing microbial community structures across AMP, ASV, and OTU pipelines based on Bray-Curtis distance.

| Comparison Pair | Mantel Statistic (r) | P-Value | Method |
| --- | --- | --- | --- |
| AMP vs ASV | 0.83 | 0.001 | Spearman |
| ASV vs OTU | 0.96 | 0.001 | Spearman |
| AMP vs OTU | 0.81 | 0.001 | Spearman |

AMP, raw amplicon sequencing data; ASV, Mitag4taxa pipeline using the denoising-based ASV approach; OTU, Mitag4taxa pipeline using the 99% similarity-based OTU clustering. All Mantel tests were performed with 999 permutations based on Bray-Curtis distance matrices.

**Figure 2.**
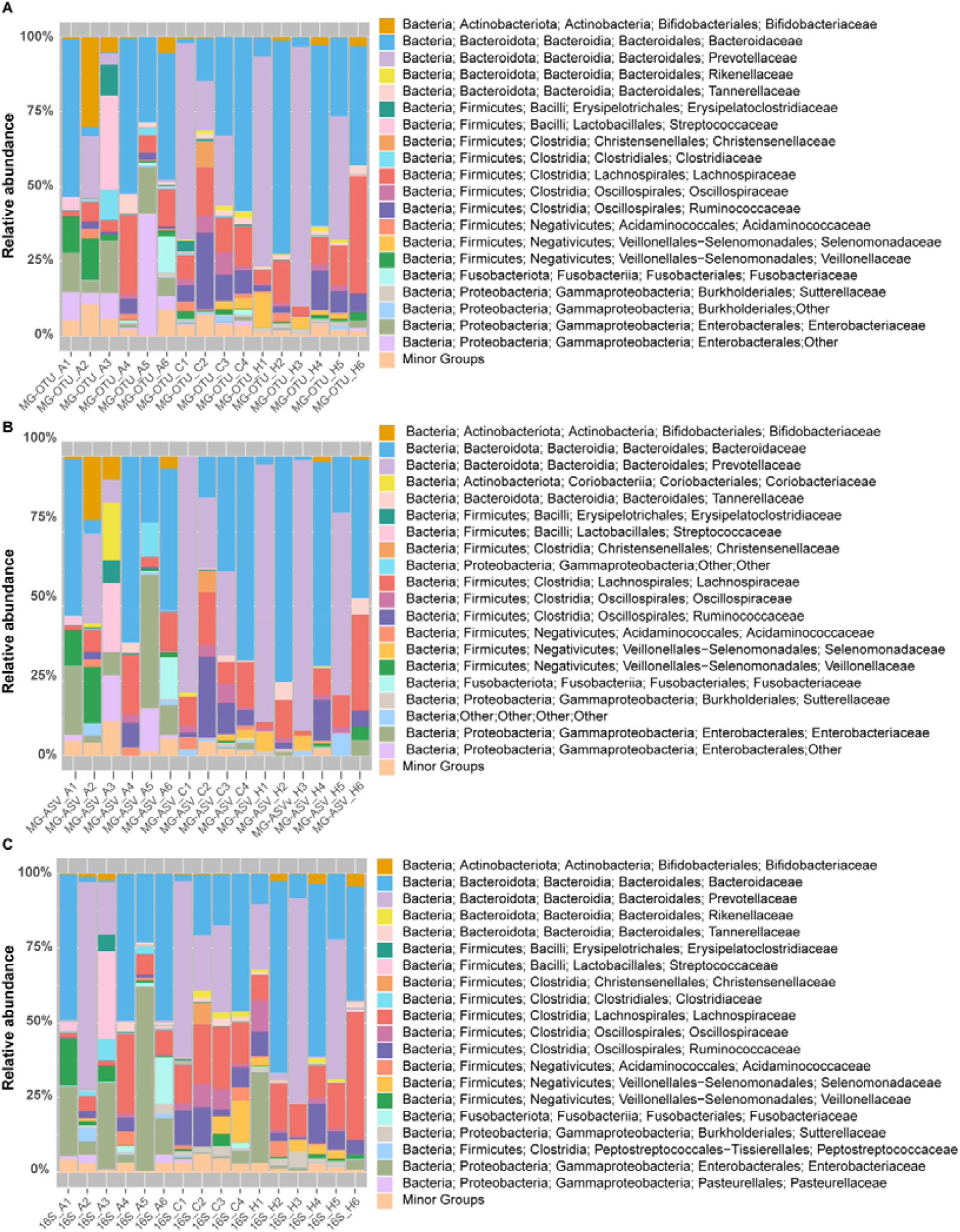
Comparison of prokaryotic composition at the family level across metagenomic and amplicon sequencing datasets. A. Relative abundance of microorganisms across samples at the family level, derived from V4 region sequences in FASTA format (without quality scores) extracted from a public metagenomic dataset using Mitag4taxa software. The OTU data (MG-OTU) derived from 16S rRNA gene V4 region were used for taxonomic classification. B. Relative abundance of microorganisms across the same samples at the family level. The V4 region sequences in FASTQ format (with quality scores) were extracted from the same metagenomic dataset using Mitag4taxa software, followed by sequence correction with DADA2. The ASV data (MG-ASV) were used for taxonomic classification. C. Relative abundance of microbial families across samples, derived from the 16S rRNA gene amplicon sequencing dataset (16S) of the same samples. All three datasets (A–C) originate from human gut microbiota samples obtained from the same set of patients (Tables S1-S3). Samples with the same identifiers (A1-6, C1-4 and H1-6) correspond to the same group of individuals across panels.

The microbial community structure derived from 16S amplicon sequences was also compared with those V4 sequence extracted from metagenomic data via Mitag4taxa to generate the MG-OTU and MG-ASV profiles. Procrustes analysis revealed a high degree of congruence between amplicon-based and metagenome-derived communities (*P* < 0.05). Notably, the MG-OTU approach (M^2^ = 0.328) showed tighter alignment with the 16S reference than the MG-ASV approach (M^2^ = 0.565) (Figure 3a and Figure 3b). Using a real dataset collected from hospital patients, we evaluated the performance of Mitag4taxa in identifying microbial community structure from metagenomic data and compared the results with those obtained from 16S rRNA amplicon sequencing. The overall taxonomic profiles derived from Mitag4taxa were highly consistent with those from amplicon sequencing, demonstrating that the software can reliably capture the dominant taxa within complex microbial communities. These results support the robustness and reliability of Mitag4taxa for accurate taxonomic profiling in microbiome research. Between the two metagenomic approaches in Mitag4taxa, the MG-ASV method performed slightly less effectively than MG-OTU. This may be attributed to limited sequencing depth in the metagenomic datasets, which affects the stringent error-correction step in DADA2. The denoising process may remove a substantial fraction of reads, leading to reduced sensitivity and underestimation of diversity. In contrast, the OTU-based approach providing a more comprehensive picture of the microbial community in cases that sequencing depth is modest.

**Figure 3.**
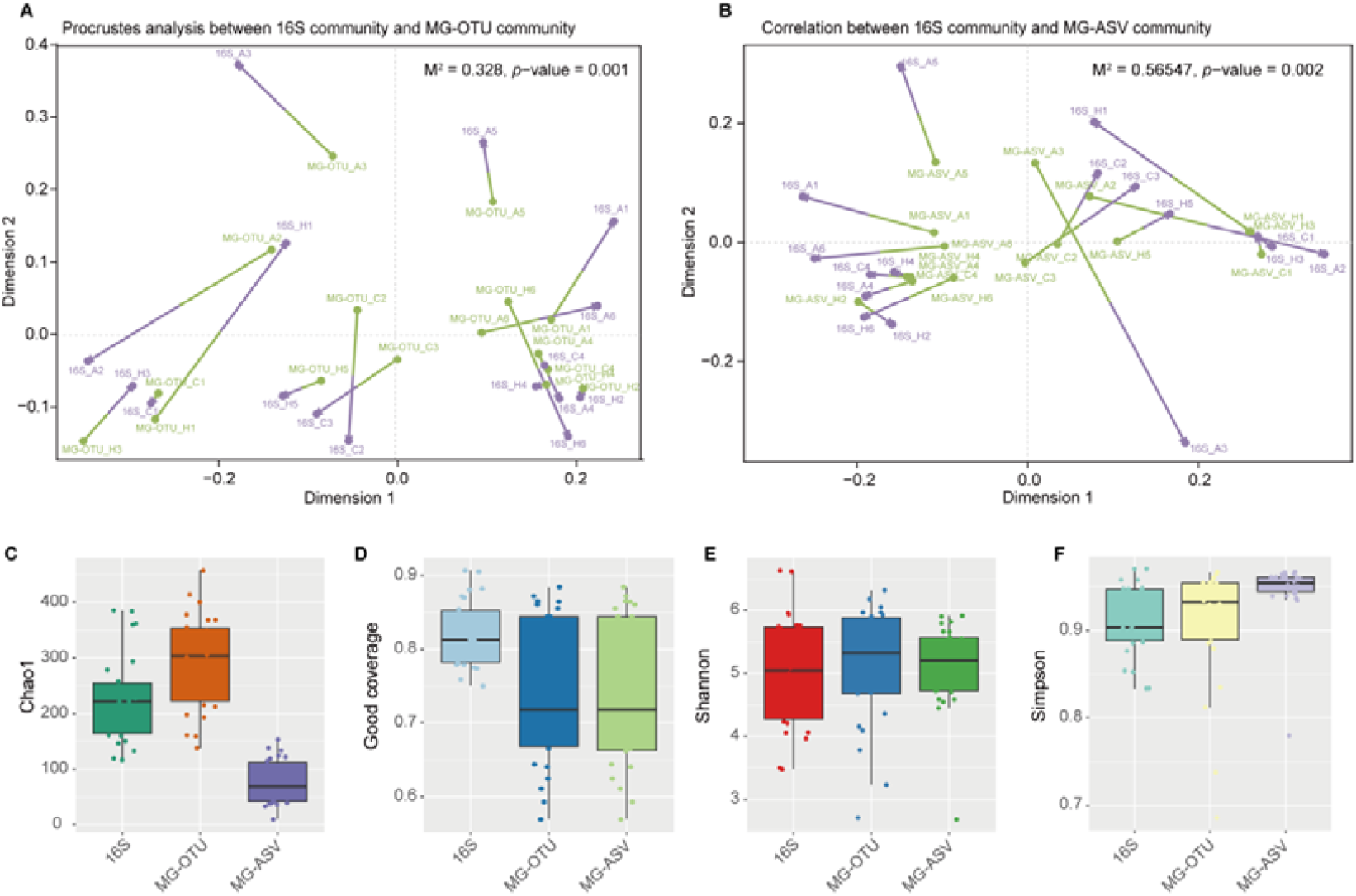
Alpha diversity and Procrustes analyses between mitag- and amplicon-based testing datasets. (A-B) Procrustes analysis evaluating community structure similarity. The plots compare the 16S rRNA gene amplicon data with mitag-derived V4 reads extracted via Mitaq4taxa OTU method (A) and extracted via Mitaq4taxa ASV method (B), respectively. Arrows connect matched samples, where purple and green represent community coordinates derived from amplicon sequencing and mitag datasets, respectively. The arrow length indicates the Procrustes residual, with shorter arrows reflecting higher structural similarity. Statistical significance is assessed via M^2^ and *P*-values. (C-F) Boxplots represent alpha diversity indices of microbial communities assessed by Chao1 (C), Good’s coverage (D), Shannon (E), and Simpson (F) across sample groups. For each index, the left boxplot generated by V4 sequences in FASTA format from the testing metagenomic dataset, while the right boxplot corresponds to the matching amplicon sequencing data.

To evaluate the efficacy of the different modules within the software, the mitag4taxa was used to extract SSU and LSU sequences from the same human gut metagenomic dataset. A comparative analysis of the resulting community structures revealed disparities in taxonomic assignment. Specifically, a substantial proportion of the LSU-based results remained unassigned (mean = 78.84%), whereas the unassigned fraction in the SSU results was negligible (0.05%) (Figure S1, Tables S4 and S5). Furthermore, while the overall community profiles derived from SSU and the extracted V4 regions were largely consistent, the V4-based analysis provided higher taxonomic resolution. In several instances, taxa that were categorized as “Other” in the SSU results were successfully resolved at the family level using the V4 sequences (Figure S1, Tables S4 and S5).

To compare the alpha diversity of the communities, four parameters were calculated, Chao1, Good’s coverage, Shannon and Simpson. Chao1 is a non-parametric estimator of species richness based on the number of observed species and the counts of singletons and doubletons^28^, making it particularly suitable for samples with many rare taxa. In our analysis, the Chao1 value obtained using the OTU-based method with metagenomic data (MG-OTU) was higher than that obtained from 16S rRNA amplicon sequencing (16S), whereas the ASV-based method with metagenomic data (MG-ASV) yielded the lowest Chao1 value among all methods (Figure 3c). Good’s Coverage reflects the proportion of the total species represented in a sample and is commonly used to assess whether sequencing depth is sufficient^29^. For Good’s Coverage, 16S datasets exhibited higher values than both MG-OTU and MG-ASV (Figure 3d). The Shannon index quantifies species diversity by considering both richness and evenness; higher values indicate greater diversity, and the index is sensitive to rare species^30^. The MG-OTU datasets displayed higher Shannon values than 16S, while 16S-based Shannon values were higher than those from MG-ASV (Figure 3e). The Simpson index measures the probability that two randomly selected individuals belong to the same species; higher values indicate dominance by a few taxa and lower overall diversity^31^, making it more sensitive to dominant species (Figure 3f). Here, 16S datasets exhibited lower Simpson values than MG-OTU, while MG-OTU values were lower than MG-ASV ones.

Alpha diversity indices provided deeper insight into the distinctions between metagenomics- and amplicon-based methods. The Chao1 index, which reflects species richness and emphasizes rare taxa, was higher in the MG-OTU datasets compared with the 16S samples. This indicates that the MG-OTU-based approach implemented in Mitag4taxa is more sensitive to low-abundance microorganisms that might be missed in amplicon sequencing due to PCR amplification bias during library construction. Theoretically, Mitag4taxa as an PCR amplification independent method, can capture all taxa in a sample if the sequencing depth is sufficiently deep.

To further evaluate performance of Mitag4taxa for eukaryotic taxonomic profiling, we analysed the shotgun metagenomic dataset for Santa Barbara Basin deep-sea sediment cores (NCBI accession PRJNA766251). The Mitag4taxa-OTU approach rather than the ASV-based method was employed, as the latter requires a higher density of extracted sequences to maintain taxonomic resolution^32^. At the family level, *Collodaria, Leotiomycetes*, and *Spumellaria* consistently ranked as the top three taxa, with mean relative abundances of 21.77%, 9.52%, and 8.08%, respectively (Figure S2, Table S6). Examining the individual layers, *Collodaria* was the most abundant taxon in the majority of samples, accounting for 12.63% at a depth of 16.45 mbsf (sample 1-2_SBBasin_16), 37.40% at 11.85 mbsf (19-20_SBBasin_11), 28.83% at 7.35 mbsf (29-30_SBBasin_7), and 29.98% at 4.35 mbsf (41-42_SBBasin_4). Conversely, the fungal class *Leotiomycetes* reached its peak dominance in the shallowest sample at 1.25 mbsf (55-56_SBBasin_1), comprising 42.41% of the community. This high fungal proportion in the upper layer likely reflects previously noted growth due to oxygen exposure during core storage rather than the original paleo-ecosystem^33^.

Taxonomic profiling of eukaryotic microorganisms in metagenomic and metatranscriptomic datasets is commonly performed using tools such as Kraken2 and MetaPhlAn^9,11^. Kraken2 classifies reads using an exact k-mer matching strategy against reference genomes, whereas MetaPhlAn relies on clade-specific marker genes derived from large collections of microbial genomes. While these approaches perform well for prokaryotic communities, their performance depends strongly on the availability of well-characterized reference genomes. For many eukaryotic microorganisms, however, genomic resources remain limited. Numerous taxa have available 18S rRNA sequences but lack complete or high-quality genome assemblies, which often leads to a substantial proportion of reads remaining unclassified when genome-based approaches are applied. The MiTag4Taxa that utilizes reference databases constructed from 18S rRNA gene sequences (particularly the V9 region), can provide complementary advantages for eukaryotic community profiling. The fact is that 18S rRNA gene sequences are more broadly documented for taxonomy than eukaryotic complete genomes in public databases. As a result, this V9-based approach may improve the detection of eukaryotic taxa in environmental sequencing datasets. However, our results of the OTU module showed that metagenomic sequencing depth for eukaryotes is usually too low.

### Evaluation of Mitag4taxa using CAMI marine datasets

We evaluated the OTU function of Mitag4taxa using ten simulated datasets from the Critical Assessment of Metagenome Interpretation datasets (CAMI)^34^. The datasets were synthesized by 155 newly sequenced marine isolate genomes and 622 public genomes from MarRef^35^. The RiboTagger^26^ software was used as the control, as it uses the similar method to obtain community structure information from the metagenomics and metatranscriptomics data. With the SSU amplicon sequencing information, the taxonomic analysis was used to explore the composition in metagenomics and metatranscriptomics data with the RiboTagger software.

The prokaryotes in CAMI marine datasets were used for the evaluation first. To compare performance of the tools, four evaluation metrics were calculated: F1 Score, Weighted UniFrac Error, Bray–Curtis Distance, and L1 Norm Error. The F1 Score, which integrates precision and recall as their harmonic mean, reflects the accuracy of classification. Its values range from 0 to 1, with higher values indicating better performance. In our tests, Mitag4taxa achieved an average F1 Score of 0.80, outperforming RiboTagger, with the score of 0.1 (Figure 4A). Weighted UniFrac quantifies phylogenetic dissimilarity between predicted and true communities^36^. Bray–Curtis Distance measures compositional differences, with values closer to 1 indicating greater divergence^37^. L1 Norm Error evaluates deviations in predicted taxon abundances, where smaller values indicate higher accuracy^27^. For all three dissimilarity/error metrics (Weighted UniFrac, Bray–Curtis Distance, and L1 Norm Error), Mitag4taxa consistently outperformed RiboTagger by yielding lower values, demonstrating higher predictive accuracy and robustness.

**Figure 4.**
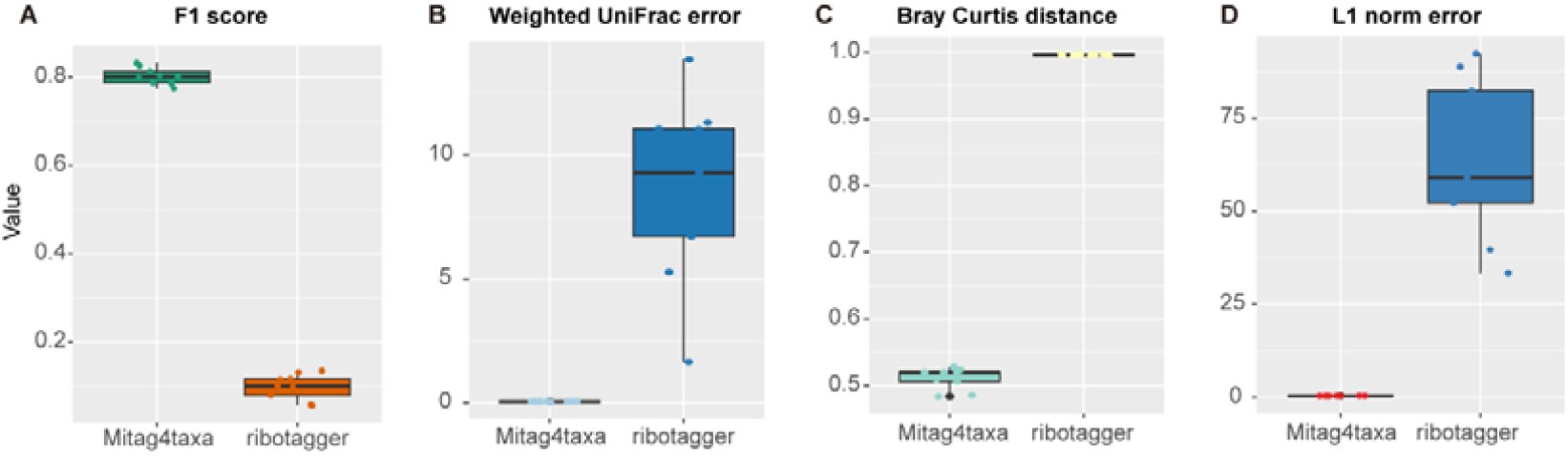
Taxonomic profiling accuracy based on CAMI marine datasets. Assessment of taxon identification accuracy (F1 score (A)) and relative abundance accuracy (Weighted UniFrac error (B), Bray Curtis distance (C), and L1 norm error (D)) at the ranks of genus, without abundance cutoff.

### Evaluation of Computational Efficiency

The benchmarking analysis reveals that mitag4taxa possesses a decisive advantage in computational efficiency when operating under constrained resource conditions. Using a fixed allocation of 2 CPU cores, mitag4taxa exhibited the shortest runtime across the six gut metagenomic samples (Figure 5). This performance gap underscores the optimized algorithmic design of mitag4taxa, which allows for rapid taxon identification fragment identification and extraction without the need for alignment of all the genome from the database.

**Figure 5.**
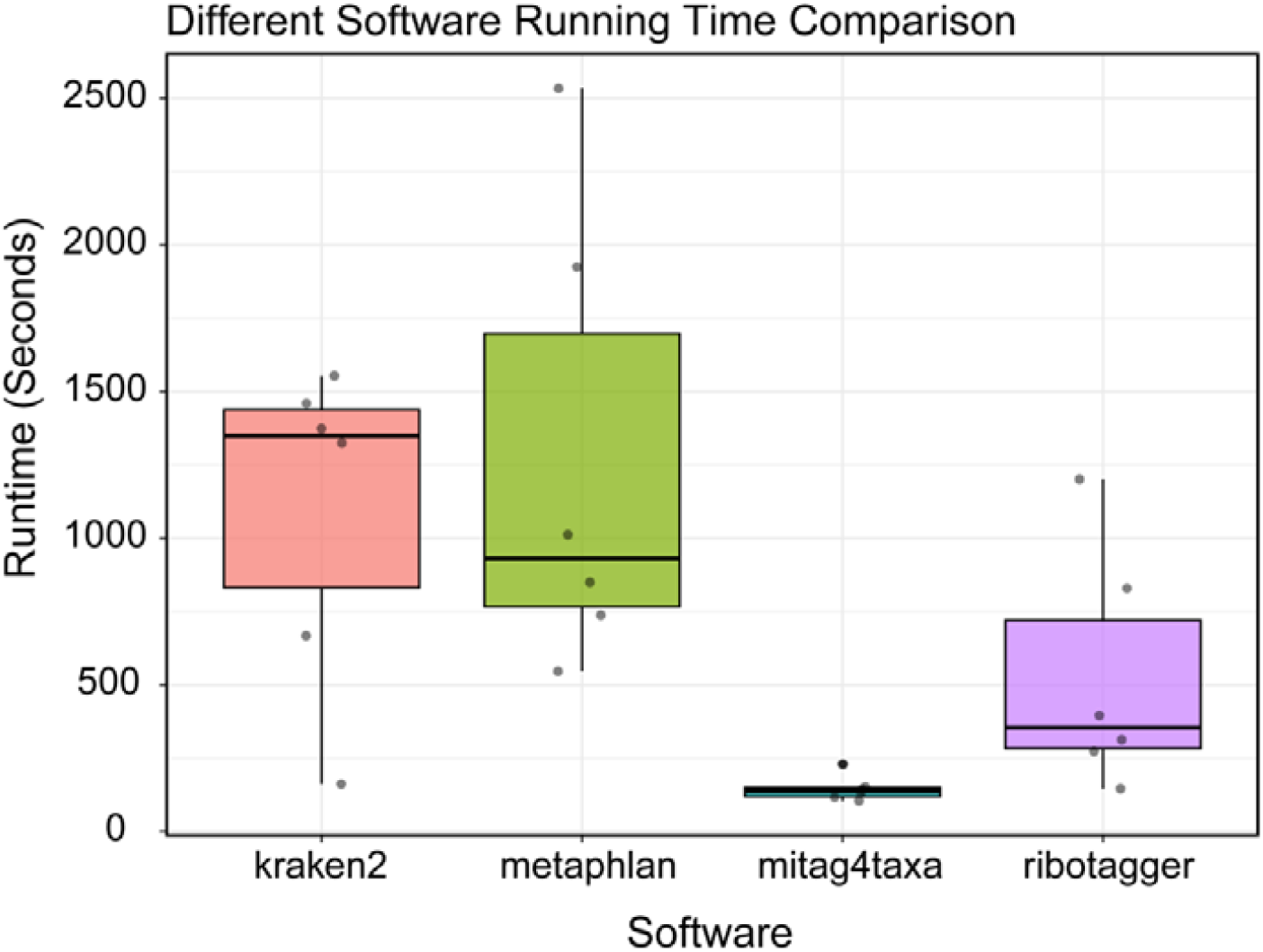
Computational efficiency benchmarking of bioinformatic pipelines. The boxplot illustrates the distribution of runtime performance for kraken2, metaphlan, mitag4taxa, and ribotagger. Benchmarking was performed using six gut metagenomic samples (A1–A6, corresponding to Figure 2) to evaluate performance across representative biological datasets. To ensure experimental rigor and comparability, all tools were executed on an identical hardware configuration with a fixed resource allocation of two cores of a 1.5-GHz Intel(R) Xeon(R) Platinum 826e CPU.

Mitag4taxa provides an efficient strategy for identify microbial community composition and estimating relative abundance from metagenome and metatranscriptome by utilizing rRNA gene sequence. By employing a regularly updated HMM database constructed from SILVA and PR2 databases, Mitag4taxa can achieve accurate taxonomic classification of microorganisms from complex environmental samples. Its HMM-based algorithm is outstanding in capturing deep evolutionary signatures, making it powerful for exploring “Microbial Dark Matter” in extreme environments (e.g., deep sea or hydrothermal vents)^38^. Unlike sequence-matching tools, Mitag4taxa can identify highly divergent or novel taxa by recognizing conserved structural patterns, ensuring high sensitivity even when reference genomes are sparse. Besides, the lightweight design and low computational requirements of the software make it particularly suitable for use in resource-limited environments, such as on personal computers, thereby expanding the accessibility of metagenomic analyses to a wider range of users and laboratories.

Despite these advantages, Mitag4taxa still faces certain limitations. The accuracy of taxonomy assignment largely relies on the completeness and quality of the reference database. Incomplete or biased reference data may limit the accurate identification of novel or poorly represented taxa, particularly in complex or extreme environments where genomic resources are still scarce. To address this challenge, continuous updating of the SSU rRNA reference database is planned, aiming to incorporate newly characterized taxa and improve taxonomic resolution across diverse microbial lineages. In addition, the evaluation of the 18S module is currently constrained by the lack of standardized benchmark datasets. Unlike prokaryotic 16S profiling, for which community benchmarking initiatives such as the CAMI framework provide well-curated validation datasets, comparable high standard datasets for eukaryotic 18S-based metagenomic profiling are still limited. This restricts the ability to perform systematic quantitative benchmarking of eukaryotic taxonomic reconstruction. Future work will therefore focus on expanding the 18S reference database and incorporating more publicly available metagenomic datasets containing well-characterized eukaryotic communities, which will facilitate more rigorous validation and improve classification robustness. Besides, future development of Mitag4taxa will focus on enhancing both its scope and artificial intelligence integrated functionality. In addition to regular database updates, forthcoming versions will incorporate modules for viral identification, thereby extending its applicability beyond prokaryotic and eukaryotic communities. Such improvements will further strengthen its ability to provide comprehensive insights into microbial ecology and clinical application from metagenomic and metatranscriptomic data.

## Contributions

**Yinghui He**: Conceptualization, Investigation, Methodology, Visualization, Data curation, Writing – original draft. **Yiling Du**: Data curation, **Loi Nguyen**: Resources, **Yong Wang**: Conceptualization, Funding acquisition, Resources, Supervision, Writing – review & editing.

## Software availability

The Mitag4taxa software, including a comprehensive user manual, installation scripts, and example datasets, is hosted on GitHub (https://github.com/heyinghui22/Mitag4taxa).

## Acknowledgments

This study was supported by the National Natural Science Foundation of China (42376149) and Shenzhen Key Laboratory of Advanced Technology for Marine Ecology (ZDSYS20230626091459009).

## References

1 Sereika, M. et al. Genome-resolved long-read sequencing expands known microbial diversity across terrestrial habitats. Nat Microbiol 10, 2018–2030 (2025).

2 Pinto, Y. & Bhatt, A. S. Sequencing-based analysis of microbiomes. Nature Reviews Genetics 25, 829–845 (2024).

3 Wang, Y. & Qian, P.-Y. Conservative fragments in bacterial 16S rRNA genes and primer design for 16S ribosomal DNA amplicons in metagenomic studies. PloS one 4, e7401 (2009).

4 Gloor, G. B. et al. Microbiome profiling by illumina sequencing of combinatorial sequence-tagged PCR products. PloS one 5, e15406 (2010).

5 Yang, B., Wang, Y. & Qian, P.-Y. Sensitivity and correlation of hypervariable regions in 16S rRNA genes in phylogenetic analysis. BMC bioinformatics 17, 135 (2016).

6 Stuart, J. et al. A comparison of two gene regions for assessing community composition of eukaryotic marine microalgae from coastal ecosystems. Scientific Reports 14, 6442 (2024).

7 Huson, D. H. et al. MEGAN community edition-interactive exploration and analysis of large-scale microbiome sequencing data. PLoS computational biology 12, e1004957 (2016).

8 Smith, R. H., Glendinning, L., Walker, A. W. & Watson, M. Investigating the impact of database choice on the accuracy of metagenomic read classification for the rumen microbiome. Animal Microbiome 4, 57 (2022).

9 Lu, J. et al. Metagenome analysis using the Kraken software suite. Nature protocols 17, 2815–2839 (2022).

10 Lu, J., Breitwieser, F. P., Thielen, P. & Salzberg, S. L. Bracken: estimating species abundance in metagenomics data. PeerJ Computer Science 3, e104 (2017).

11 Blanco-Míguez, A. et al. Extending and improving metagenomic taxonomic profiling with uncharacterized species using MetaPhlAn 4. Nature biotechnology 41, 1633–1644 (2023).

12 Duncan, A. et al. Metagenome-assembled genomes of phytoplankton microbiomes from the Arctic and Atlantic Oceans. Microbiome 10, 67 (2022).

13 He, Y., Zhang, H., Baltar, F. & Wang, Y. Transcriptional Difference of Deep-Sea Microorganisms under Different Sampling Methods. Environmental Science & Technology 59, 11653–11665 (2025).

14 He, Y., Baltar, F. & Wang, Y. Seasonal variability in community structure and metabolism of active deep-sea microorganisms. The ISME Journal 19, wraf214 (2025).

15 Finn, R. D., Clements, J. & Eddy, S. R. HMMER web server: interactive sequence similarity searching. Nucleic acids research 39, W29–W37 (2011).

16 Quast, C. et al. The SILVA ribosomal RNA gene database project: improved data processing and web-based tools. Nucleic acids research 41, D590–D596 (2012).

17 Guillou, L. et al. The Protist Ribosomal Reference database (PR2): a catalog of unicellular eukaryote small sub-unit rRNA sequences with curated taxonomy. Nucleic acids research 41, D597–D604 (2012).

18 Rozewicki, J., Li, S., Amada, K. M., Standley, D. M. & Katoh, K. MAFFT-DASH: integrated protein sequence and structural alignment. Nucleic Acids Res 47, W5– W10 (2019). 10.1093/nar/gkz342

19 Wang, Y., Tian, R. M., Gao, Z. M., Bougouffa, S. & Qian, P.-Y. Optimal eukaryotic 18S and universal 16S/18S ribosomal RNA primers and their application in a study of symbiosis. PloS one 9, e90053 (2014).

20 Chen, S., Zhou, Y., Chen, Y. & Gu, J. fastp: an ultra-fast all-in-one FASTQ preprocessor. Bioinformatics 34, i884–i890 (2018).

21 Shen, W., Le, S., Li, Y. & Hu, F. SeqKit: a cross-platform and ultrafast toolkit for FASTA/Q file manipulation. PloS one 11, e0163962 (2016).

22 Bolyen, E. et al. Reproducible, interactive, scalable and extensible microbiome data science using QIIME 2. Nature biotechnology 37, 852–857 (2019).

23 Callahan, B. J. et al. DADA2: High-resolution sample inference from Illumina amplicon data. Nature Methods 13, 581–583 (2016). 10.1038/nmeth.3869

24 Liu, Y. et al. Exploring gut microbiota in patients with colorectal disease based on 16S rRNA gene amplicon and shallow metagenomic sequencing. Frontiers in molecular biosciences 8, 703638 (2021).

25 Armbrecht, L. et al. Paleo-diatom composition from Santa Barbara Basin deep-sea sediments: a comparison of 18S-V9 and diat-rbcL metabarcoding vs shotgun metagenomics. ISME communications 1, 66 (2021).

26 Xie, C., Goi, C. L. W., Huson, D. H., Little, P. F. R. & Williams, R. B. H. RiboTagger: fast and unbiased 16S/18S profiling using whole community shotgun metagenomic or metatranscriptome surveys. BMC Bioinformatics 17, 508 (2016). 10.1186/s12859-016-1378-x

27 Meyer, F. et al. Critical assessment of metagenome interpretation: the second round of challenges. Nature methods 19, 429–440 (2022).

28 Chao, A. Nonparametric estimation of the number of classes in a population. Scandinavian Journal of statistics, 265–270 (1984).

29 Good, I. J. & Toulmin, G. H. The number of new species, and the increase in population coverage, when a sample is increased. Biometrika 43, 45–63 (1956).

30 Li, D., Liu, C.-M., Luo, R., Sadakane, K. & Lam, T.-W. MEGAHIT: an ultra-fast single-node solution for large and complex metagenomics assembly via succinct de Bruijn graph. Bioinformatics 31, 1674–1676 (2015).

31 Simpson, E. H. Measurement of diversity. nature 163, 688–688 (1949).

32 Joos, L. et al. Daring to be differential: metabarcoding analysis of soil and plant-related microbial communities using amplicon sequence variants and operational taxonomical units. BMC Genomics 21, 733 (2020). 10.1186/s12864-020-07126-4

33 Yang, Y. et al. Effects of oxygen availability on mycobenthic communities of marine coastal sediments. Scientific Reports 13, 15218 (2023). 10.1038/s41598-023-42329-1

34 Meyer, F. et al. CAMI Benchmarking Portal: online evaluation and ranking of metagenomic software. Nucleic Acids Research, gkaf369 (2025).

35 Klemetsen, T. et al. The MAR databases: development and implementation of databases specific for marine metagenomics. Nucleic acids research 46, D692–D699 (2018).

36 Lozupone, C., Lladser, M. E., Knights, D., Stombaugh, J. & Knight, R. UniFrac: an effective distance metric for microbial community comparison. The ISME journal 5, 169–172 (2011).

37 Beals, E. W. in Advances in ecological research Vol. 14 1–55 (Elsevier, 1984).

38 Wang, Y. et al. Diversity and distribution of eukaryotic microbes in and around a brine pool adjacent to the Thuwal cold seeps in the Red Sea. Frontiers in microbiology 5, 37 (2014).

