## Supplementary material for "Mitag4taxa: Extracting SSU rRNA Illumina reads from metagenomes for taxonomic classification": Supplmenty Figure

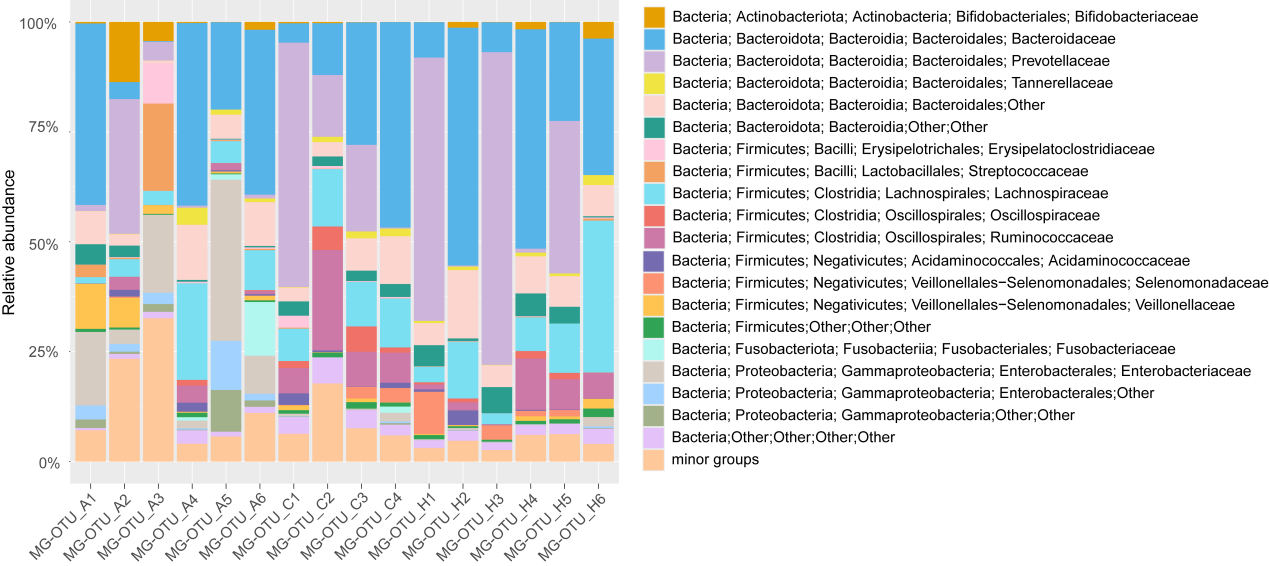

Figure S1. Taxonomic composition of prokaryotic communities based on 16S rRNA gene sequences. The stacked bar plot illustrates the relative abundance of prokaryotic taxa at the family level, derived from SSU sequences extracted by mitag4taxa (Table S5).

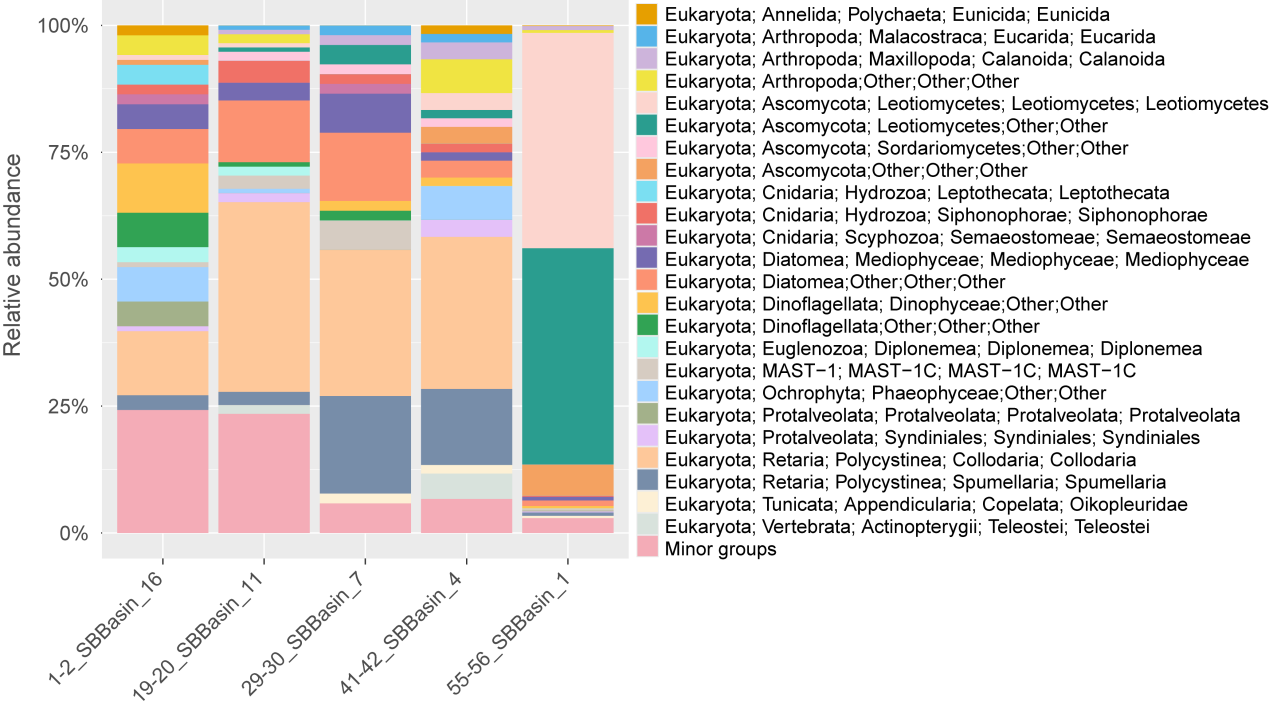

Figure S2. Relative abundance of eukaryotic microorganisms across samples at the family level, derived from V9 region sequences of 18S rRNA genes in FASTA format (without quality scores) extracted from the testing metagenomic dataset provided by NCBI using Mitag4taxa software (Table S6).
